# Synergistic and antagonistic drug interactions are prevalent but not conserved across acute myeloid leukemia cell lines

**DOI:** 10.1101/2024.02.23.581821

**Authors:** Fatma Neslihan Kalkan, Muhammed Sadik Yildiz, N. Ezgi Wood, Michael Farid, Melissa McCoy, Chengcheng Zhang, Milo Lin, Bruce Posner, Stephen S. Chung, Erdal Toprak

## Abstract

Acute myeloid leukemia (AML) is the most prevalent type of leukemia in adults. Despite advancements in medicine, the standard treatment that utilizes a combination of cytarabine and daunorubicin for AML has remained the same for decades. Combination drug therapies are proven to be an effective way to achieve targeted efficacy while minimizing drug dosage along with the unintended side effects. However, a systematic survey of synergistic potential of drug-drug interactions in the context of AML pathology is currently lacking. Here we examine the interactions between 15 frequently used cancer drugs across distinct AML cell lines and demonstrate that synergistic and antagonistic drug-drug interactions are widespread but not conserved across these cell lines. Notably, enasidenib (AG-221) and venetoclax (ABT-199), recently approved anticancer agents, exhibited the highest counts of synergistic interactions and the fewest antagonistic ones. In contrast, 6-Thioguanine (6-TG), a purine analog, was involved in the highest number of antagonistic interactions. The interactions we report here cannot be attributed solely to the inherent synergistic or antagonistic natures of these three drugs, as each drug we examined was involved in several synergistic or antagonistic interactions in the cell lines we tested. Moreover, we observed that these drug-drug interactions are not conserved across cell lines, suggesting that the success of combination therapies might vary depending on AML genotypes. For instance, we found that a single mutation in the TF1 cell line could dramatically alter drug-drug interactions, even turning synergistic interactions into antagonistic ones, as seen with AG-221 and cladribine A (2CdA). Our findings provide a preclinical survey of the potential synergistic effects revealing the complexity of the problem *in vitro*. However, the exploitable synergistic regimes in clinical scenarios remain to be explored. We anticipate these results to be an insightful guideline for future clinical studies, aiming to refine chemotherapy regimens and ultimately enhance patient outcomes.

## Introduction

Acute myeloid leukemia (AML) is a malignant blood disorder originating from clonal proliferation of abnormally or incompletely differentiated blood cells of the myeloid lineage^1^. AML is the most common leukemia type in adults^2^, affecting approximately 20,000 people in the United States with an average diagnosis age of 68^3^. Although the treatment outcomes for AML patients younger than 60 are in general better, the 5-year survival rate for patients over 65 is only about 6.9-8.9%^4^. This discrepancy is often attributed to the severe side effects associated with the intensive chemotherapy regimens that AML treatment entails^5^. This often means that aggressive treatment options are entirely off the table for older AML patients, and those with comorbidities^6^.

To address this significant clinical problem, there is growing interest in developing new drugs with fewer side effects and using combinations of existing cancer medications for greater efficacy at lower doses to reduce dose dependent side effects. For the latter, the standard approach is to look for synergistic drug combinations where combined effects of drugs are higher than the sum of each drug’s inhibitory effect when used alone^7^. However, despite several studies attempting to identify synergistic drug combinations for AML, the traditional treatment regimen for AML entailing the combined use of cytarabine and daunorubicin, remained largely unchanged for several decades^8^.

AML is a clinically and genetically heterogeneous disease^9^ which complicates the prediction of how an AML patient will respond to the cytarabine and daunorubicin combination as well as to newly developed drug(s) or drug combination therapies. This problem has become increasingly complicated and relevant with the introduction of novel drugs for AML^10^. Between 2017 and 2019, the Food and Drug Administration (FDA) approved 8 new compounds for treating AML^10^. However detailed studies assessing these compounds’ interactions with each other, or other drugs remain limited. Here, to aid clinical studies aiming to design effective combinatorial therapies for AML patients, we conducted an extensive systematic study where we measured interactions between 15 cancer drugs (see Figure 1, **Supplementary Table 4**). This drug set included commonly used anticancer drugs as well as venetoclax (ABT-199), a BCL2 inhibitor, and enasidenib (AG-221), a mutant IDH2 inhibitor. Both compounds are recently approved promising drugs designed for targeted chemotherapies^11, 12^. We tested *in vitro* efficacies of 105 drug pairs (105 = 15 x 14 / 2) against 6 standard cell lines (**Supplementary Table 5**) commonly used as an *in vitro* model for AML and other leukemias. 19 drugs were initially selected for the experiment; however, hydroxyurea, glasdegib, prednisone, and dexamethasone were excluded due to insufficient single cell drug response and killing effect. Our objectives are to find synergistic drug pairs and to determine whether these drug interactions are conserved across different cell lines.

**Figure 1.**
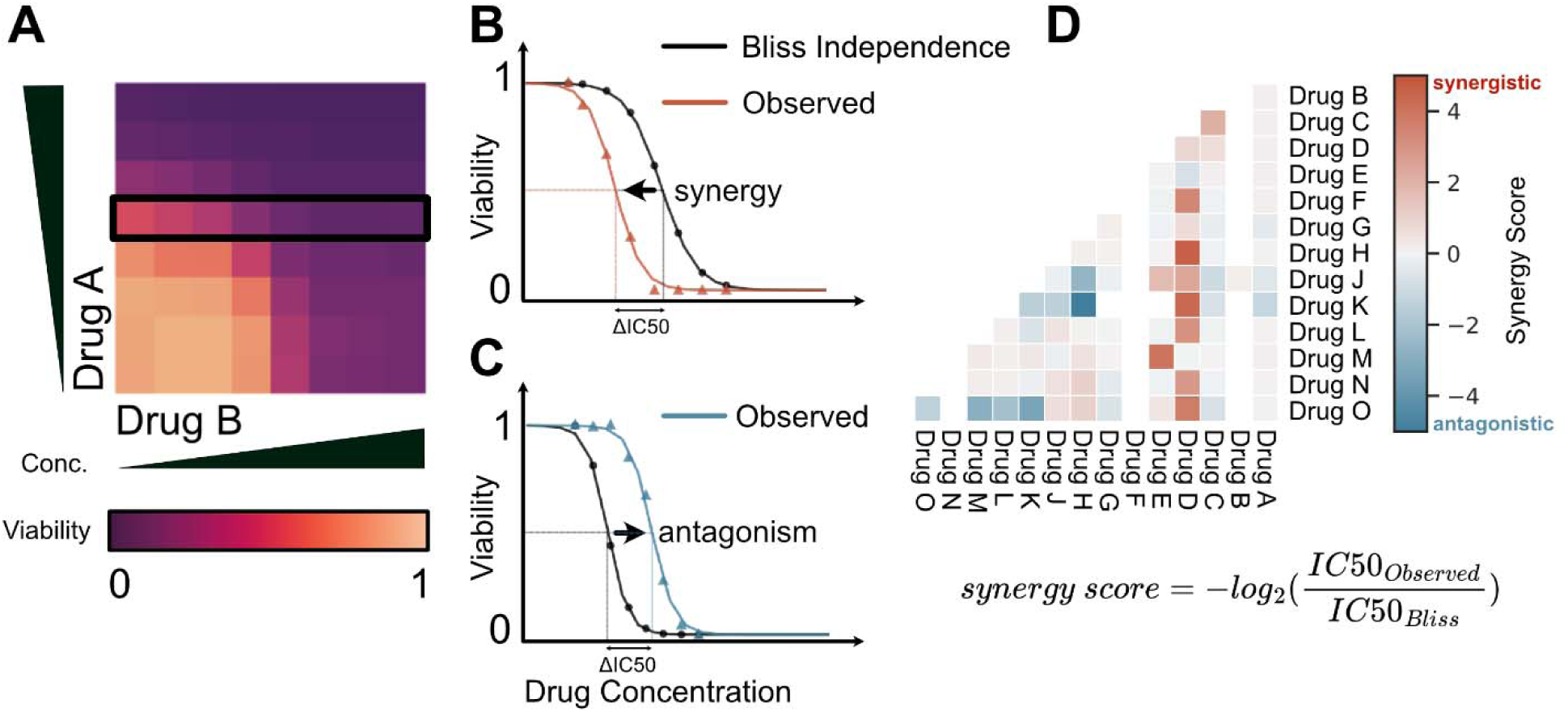
High-throughput drug combination screen. A. 105 possible drug pairs of the selected 15 drugs are screened against each other at 8 pre-selected drug concentrations. B-C. A two parameter sigmoid model (see Methods section) fitted for every row and IC50 at every specific drug range is estimated and compared with the Bliss Independence model. When the observed effect is larger than the predicted effect (panel B, red line), drug-drug interaction is considered synergistic. When the observed effect is lower than the predicted effect (panel C, blue line), drug-drug interaction is considered antagonistic. D. Synergy scores are predicted by calculating the median synergy score (S = -log_2_(ΔIC50), Methods) across all measured combination regimes. If the median value is positive, drug-drug interaction is marked as synergistic and the maximum achievable -log_2_(ΔIC50) value is used as the synergy score (red pixels). If the median value is negative, drug-drug interaction is marked as antagonistic and the minimum -log_2_(ΔIC50) value is used as the synergy score (blue pixels).

We quantified combined effects of 15 anti-cancer agents (**Supplementary Table 4**) against 6 different leukemia cell lines (FKH-1, HL60, TF-1, IDH2, Kasumi-1, K562; see **Supplementary Table 5**) using a high-throughput cell viability assay (CellTiter-Glo, Methods). Many of these drugs are clinically used to treat AML patients. Two of these drugs, ABT-199 and AG-221, are relatively new compounds and are considered promising for targeted leukemia therapies. FKH-1 was isolated from patient with Acute promyelocytic leukemia, HL60 was isolated from a patient who presented with FAB M2 AML, an aggressive variant of AML. The Kasumi-1 cell line originates from an acute myeloblastic leukemia (AML) patient. TF-1 cell line is derived from a patient with erythroleukemia, a subtype of AML. Meanwhile, IDH2 mutant-TF-1, which we will simply refer to as IDH2, is an isogenic cell line with the IDH2R140Q mutation, derived from the original TF-1 line^13^. These cell lines encompass high-risk (TP53 mutations in HL60, t(6;9) in FKH-1), low-risk (t(8;21) in Kasumi-1 and t(15;17) in HL60), and intermediate-risk (IDH mutations in TF-1 cells) genetic variants commonly observed in AML patients. In addition, we included the K562 erythroleukemia lymphoblast cell line that is derived from a patient diagnosed with chronic myelogenous leukemia as a control in our study.

## Results

### Drug-drug interactions are mapped across six diverse myeloid leukemia cell lines

We examined synergistic and antagonistic interactions between drug pairs by utilizing two-dimensional (2-D) drug gradients in 384-well plates. Every 2-D gradient included single drug dose response curves (DRCs) for the measurement of two drugs in the first column and first row of each plate. We used a two-parameter sigmoid model to represent each single DRC (up to 14 curve per drug), obtain MIC (minimum inhibitory concentration) and IC50 (the inhibitory concentration required to reduce cell viability by half) values for each cell line (**Supplementary Figure 1 and Supplementary Table 1**). In the layout depicted in **Figure 1A**, as the concentration of drug-B increases from left to right, the concentration of drug-A remains unchanged. Conversely, when the concentration of drug-A increases from bottom to top, the concentration of drug-B remains constant. At the starting point of each direction no drugs were added, and therefore at the bottom-left corner of each pattern the drug concentration is 0. For quality control, we incorporated both positive and negative controls on each plate to verify and gauge cell viability (further details are provided in the Methods section). By examining pairwise combinations of the 15 drugs on 6 distinct leukemia cell types, we determined the efficacy of 105 drug pairs at 3,167 possible combination regimes by performing 247,160 viability measurements.

We utilized the Bliss Independence Model as our neutral reference for measuring synergistic and antagonistic interactions between the drug pairs. As illustrated in **Figure 1**, for every pair of drugs, we examined cell viability at a constant dose of drug A while the concentration of drug B monotonously increased. First, we determined the anticipated cell viability under these conditions using the Bliss Independence Model (see **Methods**; local fits specific to each plate are employed). In **Figure 1B-C**, the expected cell viability is represented using black symbols at each concentration interval and a two-parameter sigmoid model shown as the black line. Subsequently, we contrasted the observed cell viability under these conditions to the expected cell viability. This analysis was repeated for each row (**Figure 1A**, black rectangle) with seven distinct constant concentrations of drug A.

For all cases, we determine the shifts in synergistic potency (Δlog2(IC50)) by computing the ratio between observed and predicted IC50s (Δlog2(IC50) = log2(IC50_predicted_ / IC50_observed_)). Finally, to minimize discrepancies and effects of idiosyncratic measurements in our analysis, for each 8 by 8 gradient for a given drug pair applied to a given cell line, we classify each interaction either as synergistic or antagonistic by calculating the median synergy score (S = -log_2_(ΔIC50), **Methods**) across all measured combination regimes. When the median value is positive, we label these interactions as generally synergistic and use the maximum achievable -log_2_(ΔIC50) value as the synergy score (as depicted by red pixels in **Figure 1d**). Conversely, when the median value is negative, we label these interactions as generally antagonistic, and use the minimum -log_2_(ΔIC50) value as the synergy score (as depicted by blue pixels in **Figure 1D**). Therefore, we evaluate the synergistic potential of any drug pair against any cell line by the highest achievable synergistic or antagonistic effect across all tested combination ranges of those two drugs.

We provided synergy scores for all drug-drug-cell line pairs that resulted in a greater than 2-fold change (median of two or three replicates) in ΔIC50 against at least one cell line in **Figure 2A**, and the complete heatmap of the estimated synergy scores organized by drug pairs in both axes is provided as **Supplementary Figure 2**. Notably, the synergy scores between drug pairs we provide here only constitute a measure of how well the combination worked with reference to underlying single drug performance, rather than a measure of general efficacy. Therefore, we provide the efficacy levels of each drug for each cell line and clinically relevant plasma levels of these drugs (vertical red lines) in **Figure 2B**. Together they can be used to identify the drugs with the highest efficacy against a specific genetic background and also the highest synergistic gains through combination therapies. We note that in real-world scenarios, drug pharmacokinetics and pharmacodynamics do not maintain steady drug dosage regimens as we did in our *in vitro* measurements as much higher doses are used in clinical settings compared to what is used in our experiments.

**Figure 2.**
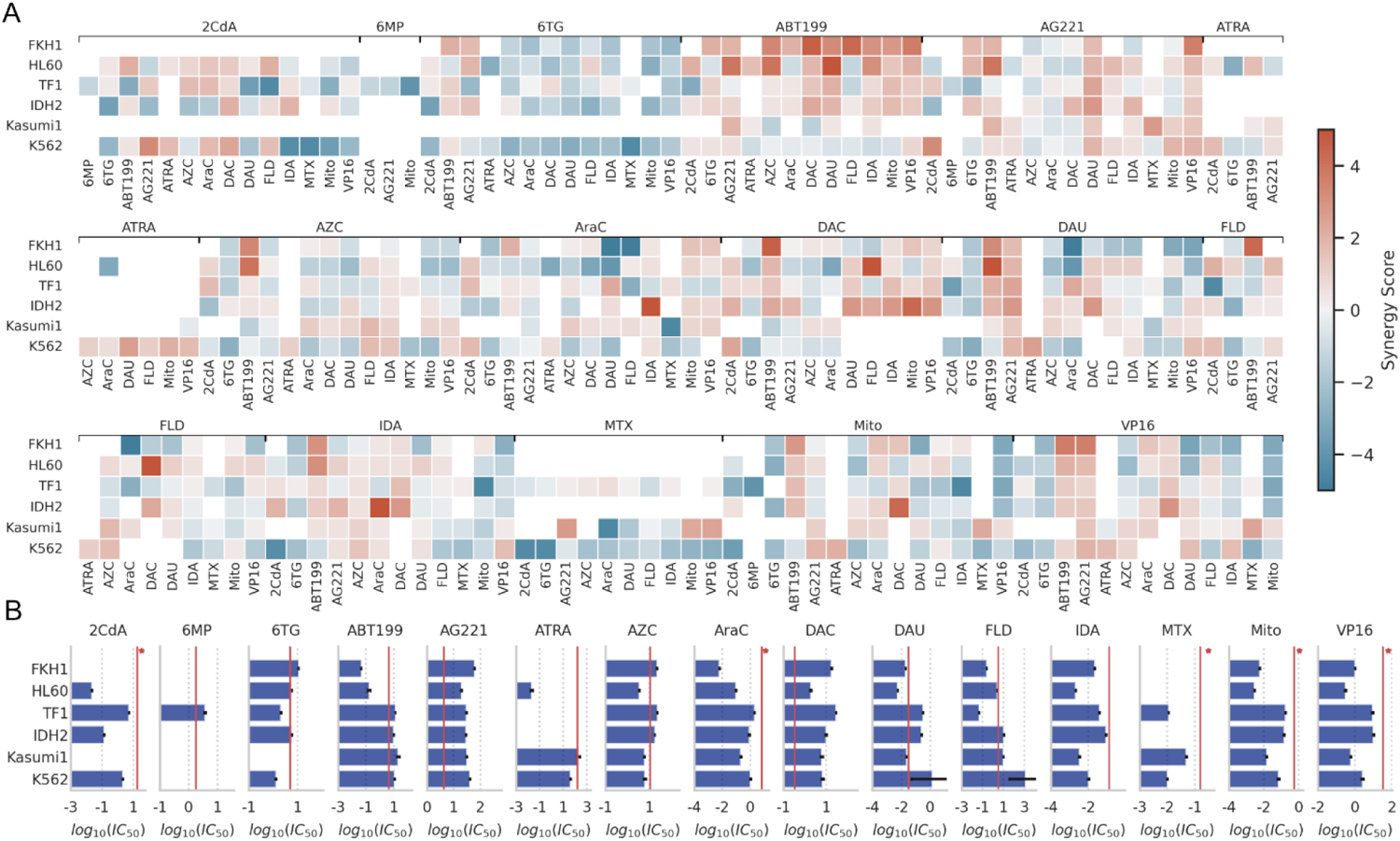
(a) Synergistic and antagonistic potency of drug pairs are revealed. Synergy scores for every drug-drug-cell line pair with greater than two-fold IC50 gain in at least one cell line are shown on the heatmap. Drug pairs are labeled at the top and bottom x-axis while cell lines are annotated at y-axis. The full interaction heatmaps are presented in **Supplementary Figure 2**. (b) IC50 values of the drugs used in this study against the six cell lines we used. Bars indicate mean values and error bars indicate standard deviation. Missing bars are due to very high IC50 values that were out of our experimental range or IC50 values that we could not unequivocally measure in our assay. Vertical red lines indicate maximum plasma levels for each drug based on previous clinical studies (**Supplementary Table 6**).

The average synergy score across the 404 evaluated drug-drug cell line pairs was - 0.005 ± 1.47 (mean ± standard deviation). This indicates that while the majority of drug interactions were independent, there was notable variability due to the large number of synergistic or antagonistic interactions. We identified 77 drug-drug-cell line pairs that had synergistic interactions, resulting in a reduction of IC50 values by at least two-fold (**Supplementary Table 3**). We identified 85 drug-drug-cell line pairs that displayed antagonistic interactions, where the combined action of the two drugs led to an increase in IC50 values by at least two-fold.

To ensure the experimental reproducibility we repeated measurements for a subset of the drug-drug cell line pairs (**Supplementary Figures 3 and 4**). For enhanced precision in these control experiments, we employed 11 by 11 drug gradients (in duplicates). We found that our measurements with 8 by 8 and 11 by 11 gradients were highly correlated (**Supplementary Figure 5**, Pearson’s correlation coefficient r = 0.67 with p = 6.7 x 10^-8^) ensuring experimental and computational reproducibility despite the significant alteration to the technical conditions of the experiment including tested drug concentration levels and intervals. Numerical values for all synergy scores and significant synergy scores can be found in **Supplementary Tables 4 and 5**, respectively.

### ABT-199 and AG-221 are all generally synergistic with most other drugs

In our study, every drug we examined exhibited synergistic interactions with at least five other drugs (**Figure 2A**). This implies that synergistic interactions are common and might not be attributed to any specific drugs. Notably, out of the significant synergistic interactions we observed, 47 out of 77 involved either ABT-199 or AG-221, which are both new generation cancer drugs. Both ABT-199 and AG-221 were involved in antagonistic interactions in a total of only 6 cases. Furthermore, these two drugs demonstrated synergistic interactions with one another across three distinct cell lines. This suggests that ABT-199 and AG-221 could potentially serve as general enhancers in AML treatments. Additionally, all six cell lines we tested had at least 8 synergistic drug-drug interactions with at least 2-fold IC50 reduction (FKH1: 14, HL60: 21, IDH2: 13, K562: 12, Kasumi1: 8, TF1: 9). These findings collectively indicate that significant synergistic drug pairs for AML treatment are likely to be identified across various genetic contexts.

### 6-Thioguanine is generally antagonistic with most other drugs

Antagonistic interactions between drugs were also widespread across all cell lines and every drug we tested (**Figure 2A**). Notably, of these interactions, 29 out of 85 involved 6-Thioguanine (6-TG), a purine analog. This suggests that in certain therapeutic combinations, particularly depending on the genetic makeup of the disease, it might be necessary to exclude 6-TG. Interestingly, the two drugs, ABT-199 and AG-221, which were highlighted for their strong general synergistic potential, showed the fewest antagonistic interactions compared to the other drugs we examined and intriguingly they also evaded the strong general antagonistic effect of 6-TG).

### Cytarabine and daunorubicin are generally independent

The “3 + 7 regimen” (3 days of daunorubicin + 7 days of cytarabine) established in the 1970s became the standard AML treatment^14^. CPX-351, a nanoliposome encapsulating cytarabine and daunorubicin in a 5:1 molar ratio, has been shown to significantly improve survival, increase complete remission rates, and facilitate more allogeneic SCTs, extending post-SCT survival. This led to its FDA approval as the primary treatment for secondary AML. This drug pair did not exhibit pronounced synergistic interactions in our experiments (**Figure 2**). Their joint effects displayed weak synergy in TF-1, Kasumi-1, and IDH2 cell lines but were weakly antagonistic in FKH-1, HL60, and K562. In view of the critical impact and characteristics of the AraC and DAU combination, we examined how the effect varies in a dose-dependent manner on each cell line and found that drugs acting independently except for the FKH-1 where we observed moderate synergy (**Supplementary Figure 8**).

### Cytarabine with alternative anthracyclines

Combining cytarabine with alternative anthracyclines such as mitoxantrone and idarubicin and establishing their optimal doses were the subject of several randomized trials^15^. Both drugs are known to be active against leukemia cell lines that are resistant to daunorubicin^16^. Whether these drugs exert a differential activity on normal hematopoietic stem cells remains unclear. However, the comparable toxicity of the three drugs in combination with conventional-dose cytarabine and etoposide during induction does not necessarily imply that the same doses of the drugs have equivalent effects with the combination of intermediate-dose cytarabine during post remission chemotherapy. In our study, we observed significant antagonism between cytarabine and daunorubicin in FKH-1 and HL60 cell lines which were partially or fully reversed when combined with idarubicin and mitoxantrone (**Figure 2**). The reverse of this shift in synergistic interaction is observed in TF1 cell line corroborating with the hypothesis that in clinical cases where the typical AraC-DAU combination is not effective, using a different anthracycline could prove beneficial.

### Conservation of drug-drug interactions is dependent on the genetic background

We assessed the conservation of drug-drug interactions across various cell lines by analyzing the correlation between synergy scores in these cell lines (**Figure 3**). The average correlation across the cell lines was 0.198 ± 0.175 (mean ± standard deviation). Considering that all varying technical conditions of the experiments including the study personnel, drug concentration intervals, culturing of the cell lines, experiment dates and randomized well selections across two experiments with duplicates our predicted synergy scores produce a Pearson correlation of 0.67 (**Supplementary Figure 5**), this indicates that the conservation of drug-drug interactions across different genetic backgrounds was weak. Among the cell lines we tested, the synergy scores from the Kasumi-1 cell line showed the weakest correlation with the synergy scores obtained from other cell lines, with an average of -0.001 ± 0.099 (**Supplementary Figure 6**). In contrast, the strongest correlation was observed between the HL60 and FKH1 cell lines, with a Pearson correlation of 0.47.

**Figure 3.**
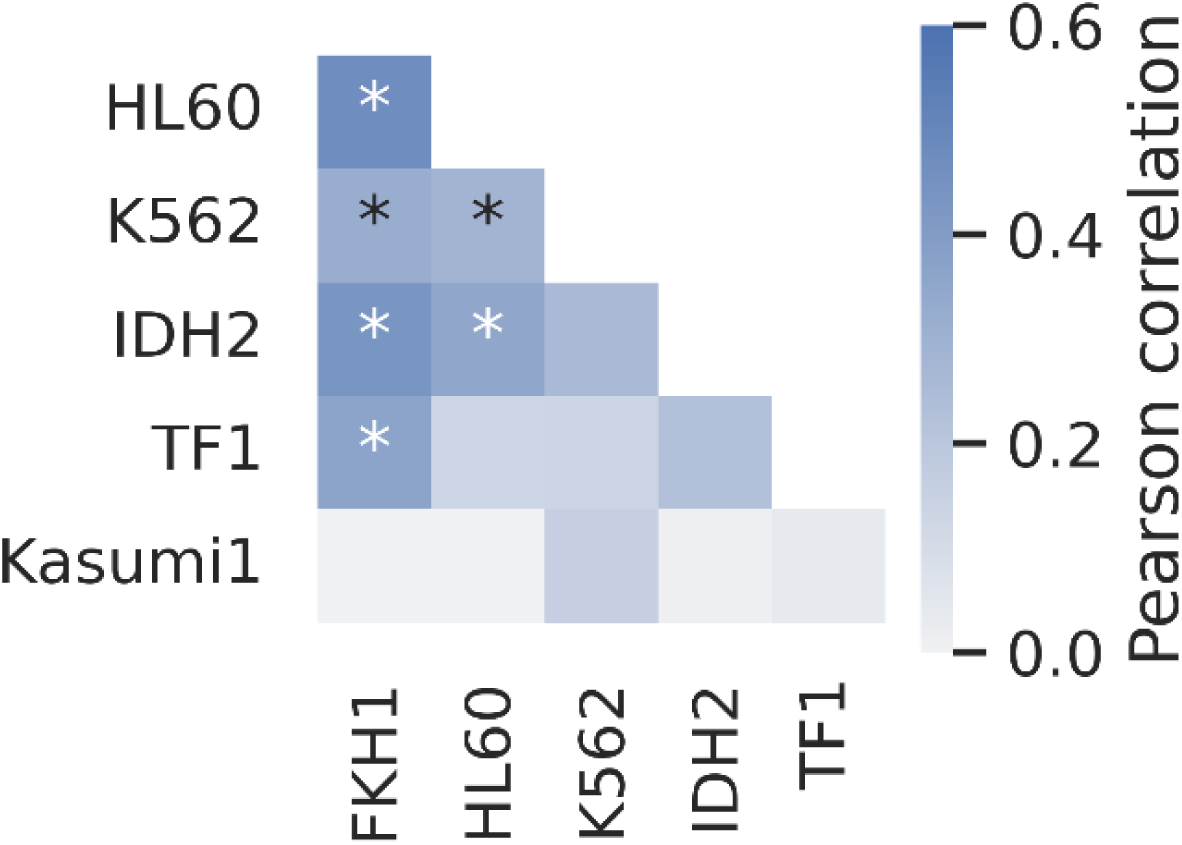
Synergistic and antagonistic drug-drug interactions are weakly correlated across cell lines. Pearson correlation between synergy scores in all cell lines are compared pairwise. Pearson correlation coefficient is colored with blue as indicated by the color bar, and star indicate where p-value is less than 0.05.

### One critical mutation can significantly alter single-drug efficacy and drug-drug interactions

Mapping variations in drug-drug interactions to the genotypes of cell lines is a complex task and beyond the scope of this study. To explore the sensitivity of this clinically important phenotype to genetic variations, we contrasted the synergy scores for drug pairs between the TF-1 and IDH2 cell lines. The IDH2 cell line is an isogenic derivative of the original TF-1 and carries only the IDH2-R140Q mutation^13^. IDH2, an enzyme which catalyzes α-ketoglutarate, and when mutated is known to induce changes in methylation^17^. Due to the small genetic distance between these two cell lines compared to other cells, one would expect that drug-drug interactions in IDH2 and TF-1 would be similar. However, intriguingly, the correlation of synergy scores between the TF-1 and IDH2 cell lines were significantly altered (**Figure 4**), indicating that a single critical genetic mutation can potentially shift the outcomes of combined drug treatments. Several of the additive interactions in TF-1 shifted to antagonism or synergy in the IDH2 cell line. Most strikingly, the synergistic interaction between AG-221 and 2CdA in TF-1 became antagonistic in IDH2 cells. **Figure 4** also shows examples of conserved antagonism (for instance, between mitomycin and VP16) and synergy (such as ABT-199 and DAC) in both IDH2 and TF-1 cell lines. Nonetheless, there were numerous drug-drug interactions that displayed synergy or antagonism in the TF-1 cell line but showed additive interactions in the IDH2 cell line.

**Figure 4.**
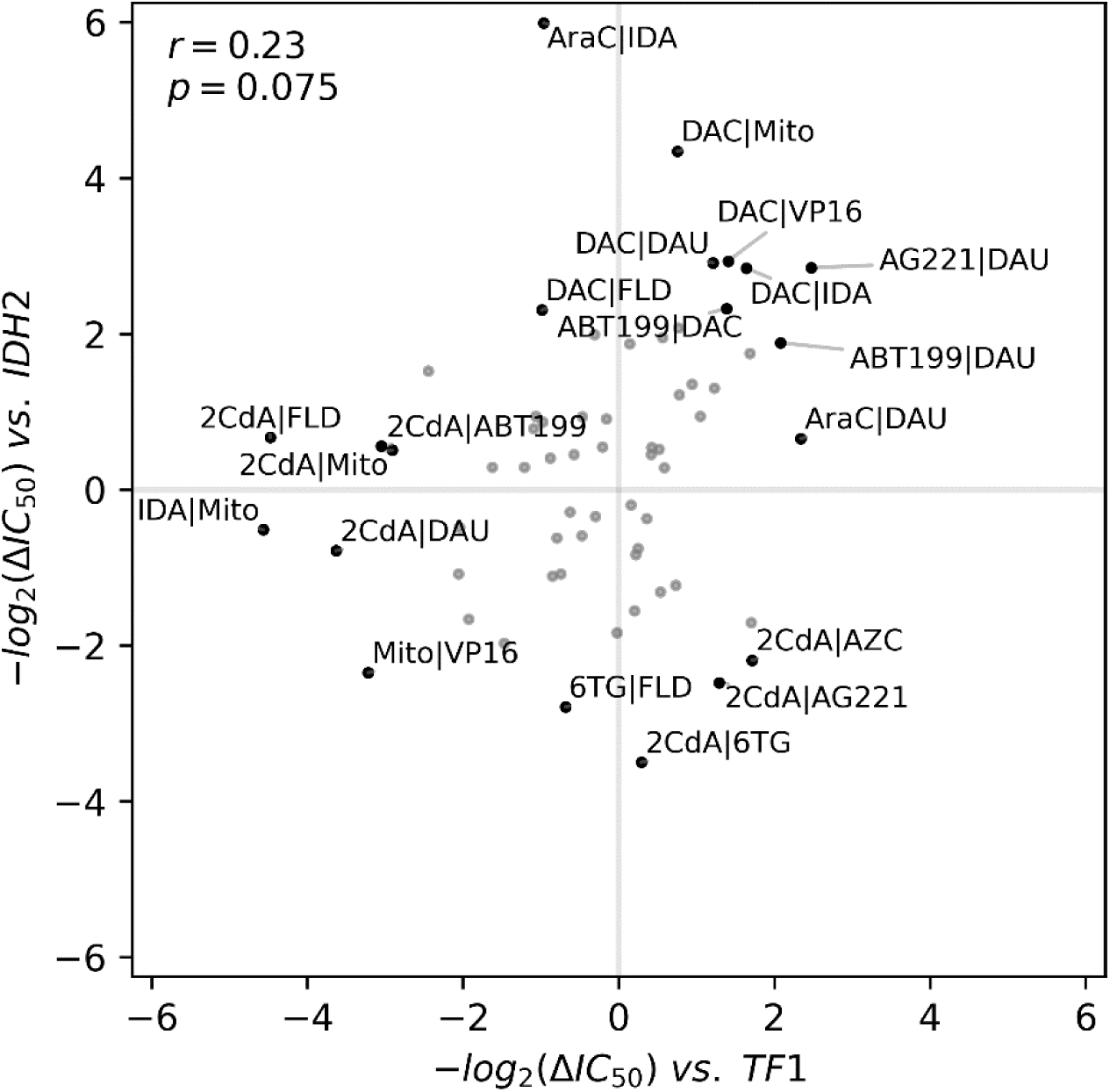
One single critical IDH2 mutation significantly alters the drug-drug interaction landscape in TF1. Measured synergy scores in cell lines IDH2 and TF1 are plotted. Pearson correlation and corresponding p-value is provided at the top left.

### Summary

In our analysis of pairwise interactions of 15 commonly used anticancer drugs in six diverse cell lines originated from myeloid leukemia, we identified numerous synergistic and antagonistic interactions between drug pairs. Notably, AG-221 and ABT-199, recent additions to the cancer treatment arsenal, accounted for the highest counts of synergistic interactions and the fewest counts of antagonistic interactions. This finding underscores the potential of both AG-221 and ABT-199 to act as synergistic agents in multi-drug therapies. Conversely, 6-Thioguanine (6-TG), a purine analog, was frequently involved in antagonistic interactions, which suggests that its utility in combination therapies for AML treatment might be limited. However, despite these noticeable interactions, it was not possible to attribute synergy or antagonism in combinatorial drug interactions to a particular drug or group of drugs, since every compound we tested was involved in several synergistic or antagonistic interactions as summarized in **Figure 2**.

We observed both synergistic and antagonistic drug-drug interactions in all cell lines we tested. However, these interactions were not universally conserved across different cell lines, as depicted in **Figure 3**. This variability suggests that the efficacy of combined therapies might be closely linked to the genetic context of the disease. As we demonstrated in **Figure 4**, even a single mutation in the TF1 cell line can profoundly alter drug-drug interactions, even converting synergistic ones to antagonistic, as seen with AG-221 and 2CdA.

In conclusion, both synergistic and antagonistic drug interactions are prevalent among anti-cancer drugs in various genetic backgrounds. Recognizing and accounting for these interactions constitutes a largely untapped resource for designing effective combination therapies for AML treatment. The findings and conclusions of this study are derived from high-throughput *in vitro* cell viability measurements, and we make no claims regarding the clinical efficacy or relevance of drug pairs. Although most of the drug regimens we use in our study are within the clinically relevant drug concentration windows in plasma (**Figure 2B**), it remains uncertain whether our observations will hold true in preclinical and clinical settings. Nevertheless, our research can offer valuable insights and serve as a foundation for clinical studies, potentially enhancing the efficacy of chemotherapies and, in the long run, improving patient treatment outcomes.

## Methods

### Cell growth conditions and High-throughput assay

Cell lines were plated at the following densities in a total volume of 60 uL per well in 384 well microtiter plates (Greiner catalog 781098): HL-60 and K562 cells at 1200 cells/well, Kasumi-1 at 1600 cells/well, TFI-IDH1 mutant at 2500 cells/well and FKH01 at 10,000 cells/well. Cells were incubated overnight at 37LC and 5% CO_2_. The following day, test compounds were dissolved in DMSO or aqueous solvent and added to 384-well plates using an Echo555 or Echo 655 liquid handler (Beckman, Inc). For combination assays, pairs of compounds were dosed in 8x8 grids at concentrations determined by the IC_50_ for each compound. One compound of each pair was dosed in columns and the other in rows. The effect of each compound alone was determined by assaying toxicity in the absence of the other compound (single agent control). All compounds were tested against themselves to measure additive effects (sham control). DMSO treated wells were included as vehicle controls for normalization. After compound addition, cells were incubated for 96 hours at 37LC and 5% CO_2_. Following the incubation period, we added 10μL Cell Titer Glo reagent diluted 1:2 (Promega, Inc.) to each well and mixed. Plates were incubated for 5 min at room temperature, and luminescence was measured using an Envision multimodal plate reader (PerkinElmer, Inc.). Relative luminescence units were normalized to DMSO (vehicle) wells.

### Data analysis

Every measurement was done in either duplicates or triplicates for every drug pair, drug concentration, and cell line. Raw absorbance values are normalized by using the median of at least 8 negative and inhibitor control wells on the same 384-plate. Equation 1 illustrates this step:

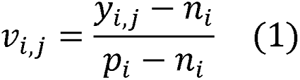

where *V_i,j_* is normalized viability, *y_i,j_* is measured absolute absorbance intensities, *n_i_* is the median of inhibitor controls, and *p_i_* is the median of positive controls. The median of all replicates of normalized viability measurements are used to create a single 8-by-8 or 11-by-11 matrix per drug pair per cell line, where each row represents increasing concentration gradient for drug A, and each column for drug B (**see Figure 1**). The first row and the first column of these raw viability matrices represent single drug response curves (**Supplementary Figure 1**). The Bliss independence model was calculated by multiplying individual effects of each drug at each concentration interval and used as the reference baseline. A two-parameter sigmoid model was fitted for estimating IC_50_ for drug A at every drug B level in logarithmic scale with a base of 2. Equation 2 is used for estimating two parameters *b_pos_* representing the IC_50_ position, and *b_shape_* representing the steepness of the dose response:

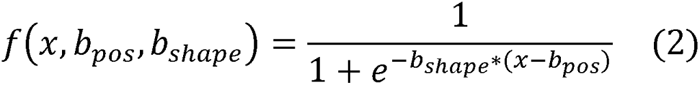

where *x* is the base 2 logarithm of the drug concentration. For achieving continuity in the model fitting, zero drug level is approximated by a 2-fold decrease from the next lowest drug concentration level. Model optimization was done by solving the non-linear least squares problem using the Levenberg–Marquardt algorithm. Synergy or antagonism of a specific drug pair and cell line combination is quantified as follows:

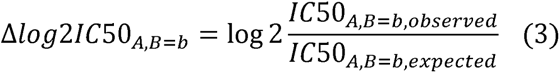

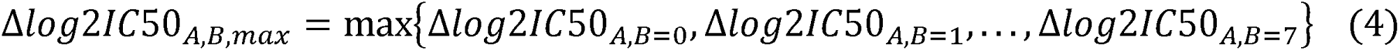

where A is the anchor drug A, and B is the library drug B, *b* is the index for increasing drug concentrations of the drug B. The direction of the synergy or antagonism is defined by the sign of the median IC_50_ change at all concentrations per drug pair cell line combination. For easier readability the synergy score is defined as *-log_2_(Δ1C50_A,B,max_)* and used as a metric for measuring synergism in the figures where synergy is positive, and antagonism is negative, and the base score of 1 represents a 2-fold change in IC_50_.

## Funding

This work is supported by UTSW Endowed Scholars Program, Human Frontiers Science Program Research grant RGP0042/2013, DOD PR172118, and Welch Foundation I-2082-20210327.

## Author Contributions

E.T., F.N.K., C.C.Z., B.P. and S.S.C. designed the research; F.N.K., and M.M. performed experiments. M.S.Y., E.T., N.E.W., M.F., and F.N.K. analyzed data. M.S.Y., E.T., and F.N.K. wrote the manuscript. All authors revised the manuscript and approved its final version.

## Declaration of Interests

E.T. is a co-founder of BAIT-BIO LLC. At the time of this publication N.E.W. is employee of Bristol Myers Squibb.

---

**Supplementary Figure 1.**
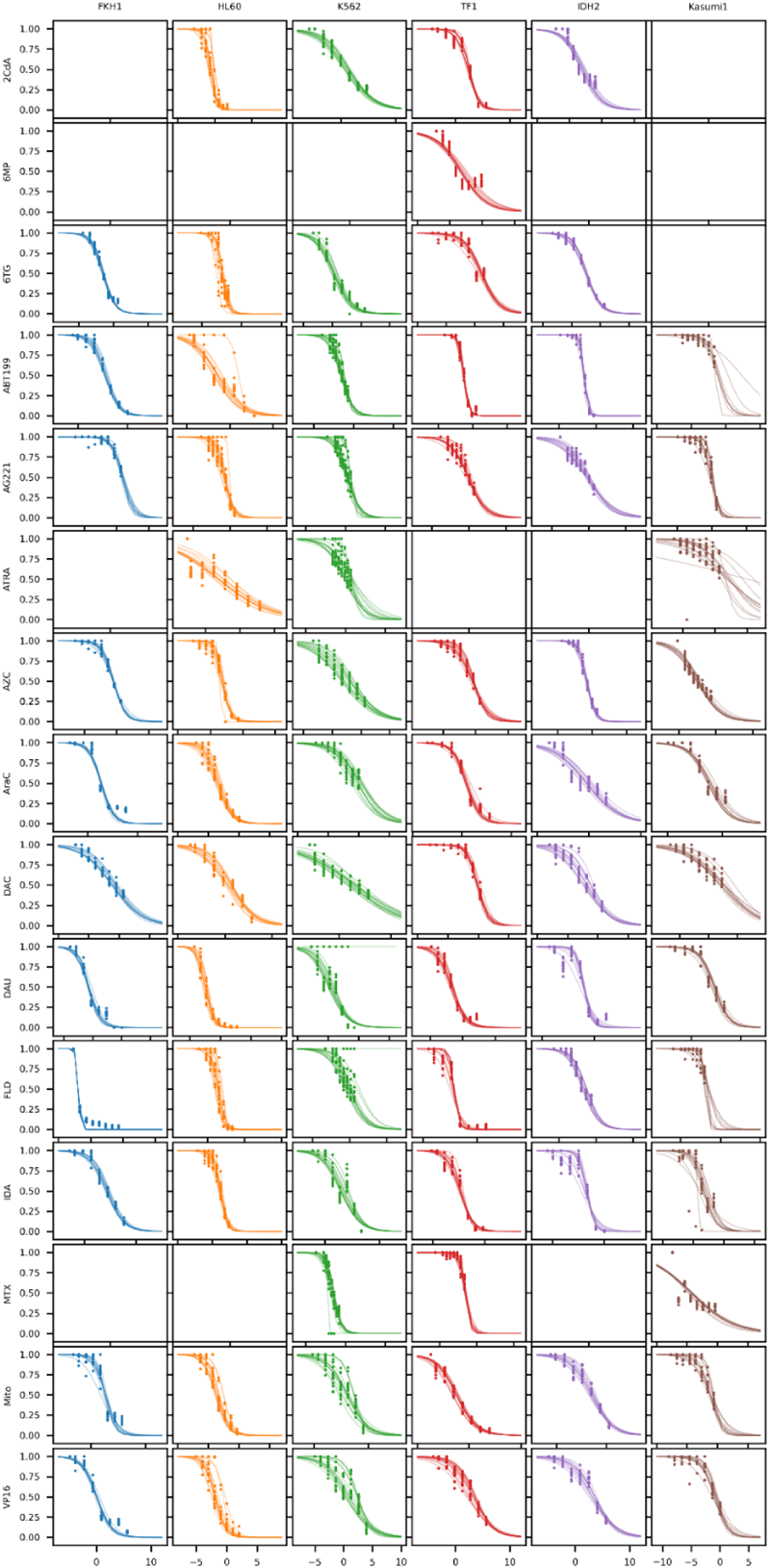
Single drug response curves measured for all drugs vs. all cell lines. For every drug and cell line pair there are 14 curves coming from the first rows and columns of the raw viability matrices. Markers in the plot show measure viability, and lines show two-parameter sigmoid model fits. Some of the drug cell line pairs are omitted due to insufficient killing by the single drug agent.

**Supplementary Figure 2.**
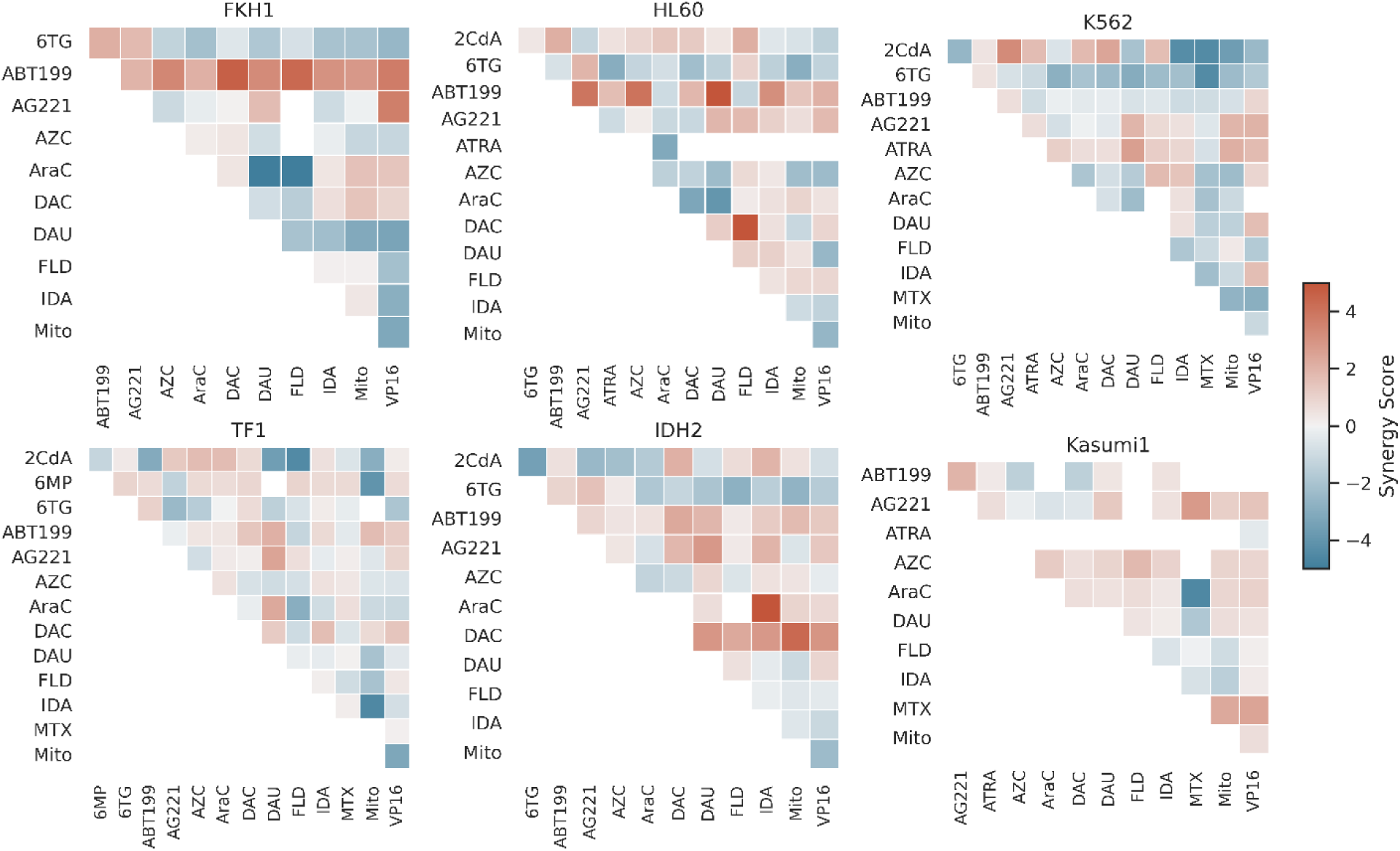
Synergy scores grouped by cell lines. All estimated synergy score included here except where either a) single drug response curves did not achieve sufficient killing, b) observed drug combination did not achieve sufficient killing, c) two parameter sigmoid model failed to fit.

**Supplementary Figure 3.**
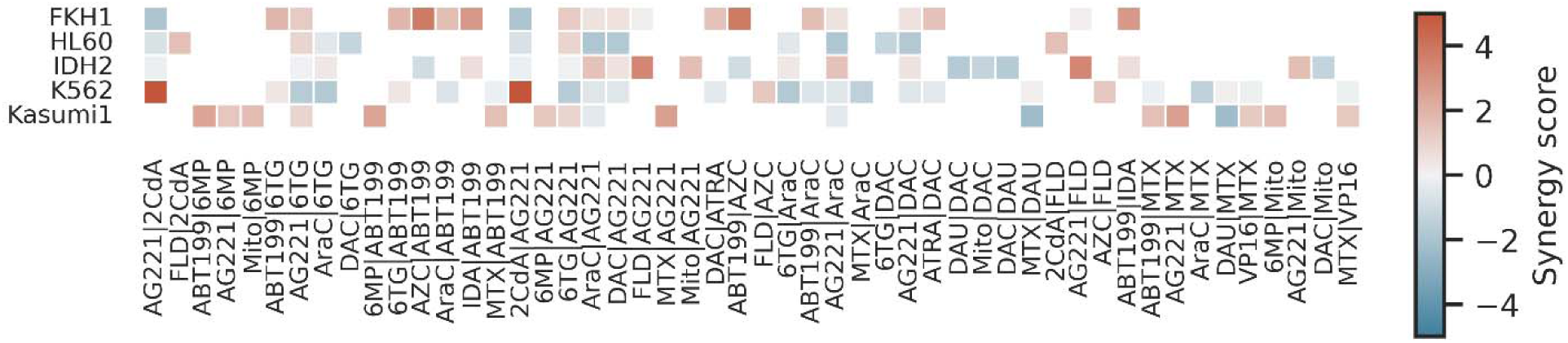
Synergy scores measured by the 11-by-11 experiment. Only drug pairs that showed greater than two-fold change in IC50 in at least one cell line included. No color shown where either a) single drug response curves did not achieve sufficient killing, b) observed drug combination did not achieve sufficient killing, c) two parameter sigmoid model failed to fit.

**Supplementary Figure 4.**
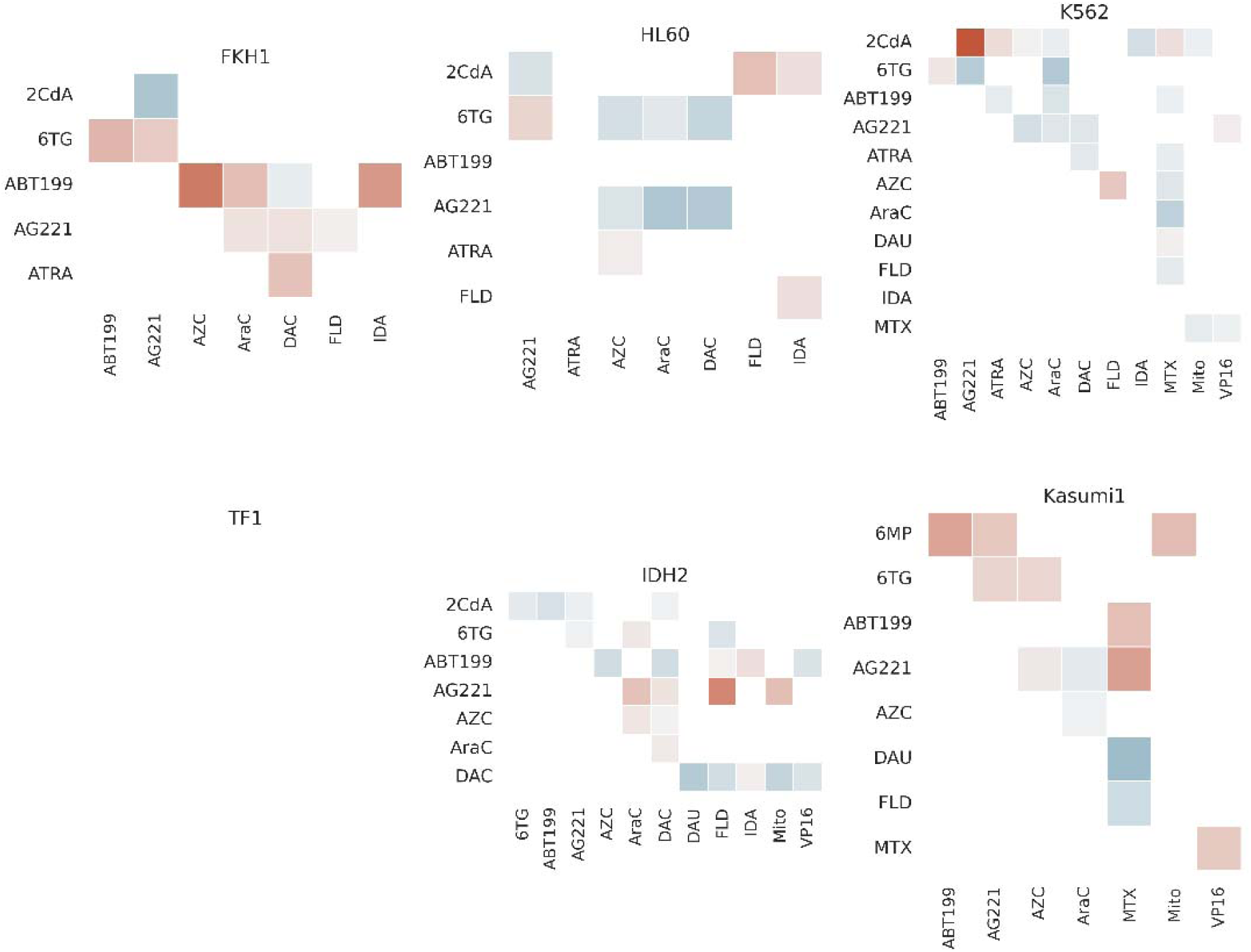
Synergy scores measured by the 11-by-11 experiment grouped by cell lines. All estimated synergy scores included here except where either a) single drug response curves did not achieve sufficient killing, b) observed drug combination did not achieve sufficient killing, c) two parameter sigmoid model failed to fit. TF1 cell line was not included in thi experiment.

**Supplementary Figure 5.**
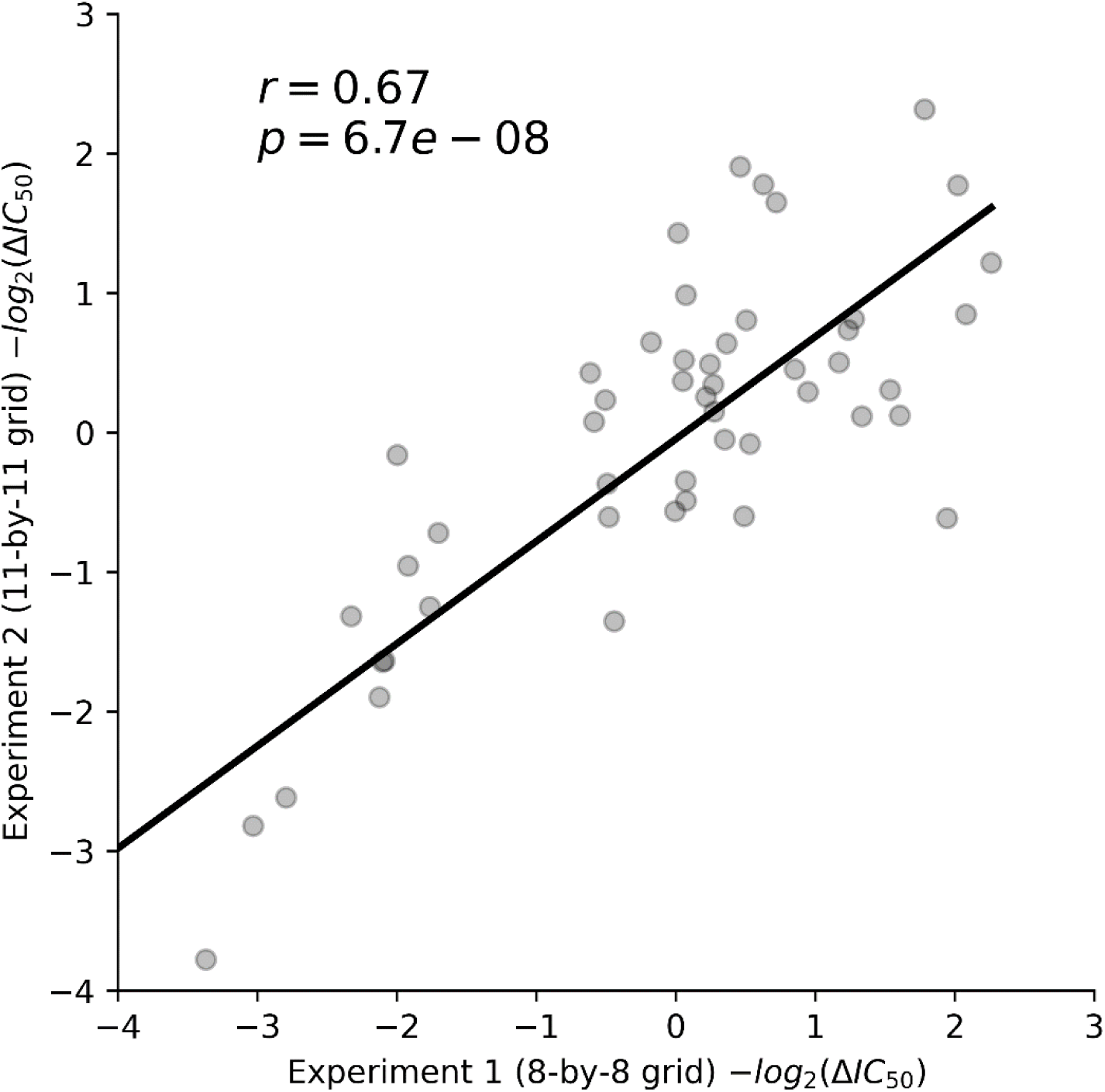
Correlation between the 8-by-8 experiment and the 11-by-11 experiment. Here we include all drug-drug-cell line pairs that produced a synergy score estimate in both experiments.

**Supplementary Figure 6.**
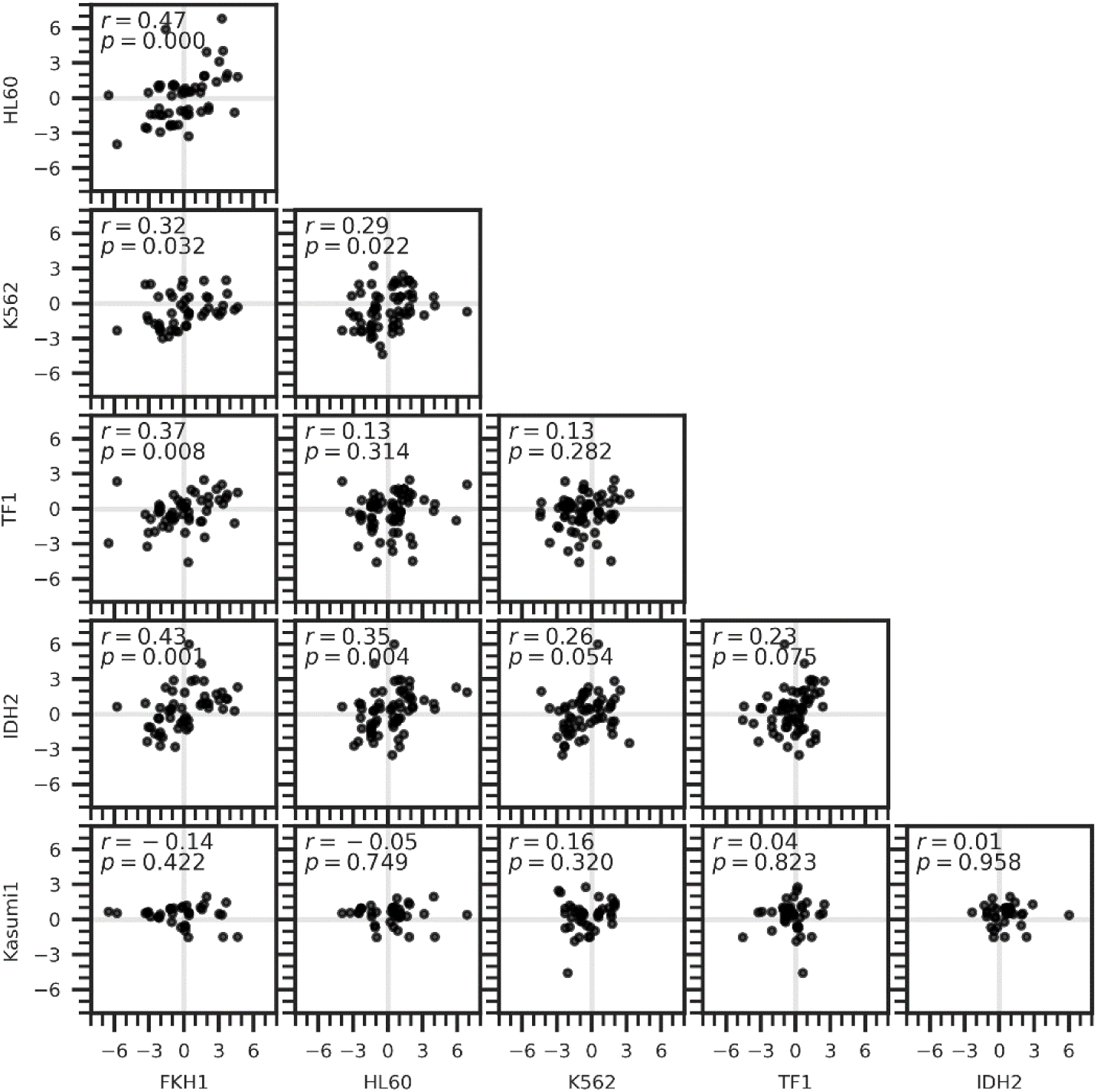
Correlation between synergy scores between all cell line pairs.

**Supplementary Figure 7.**
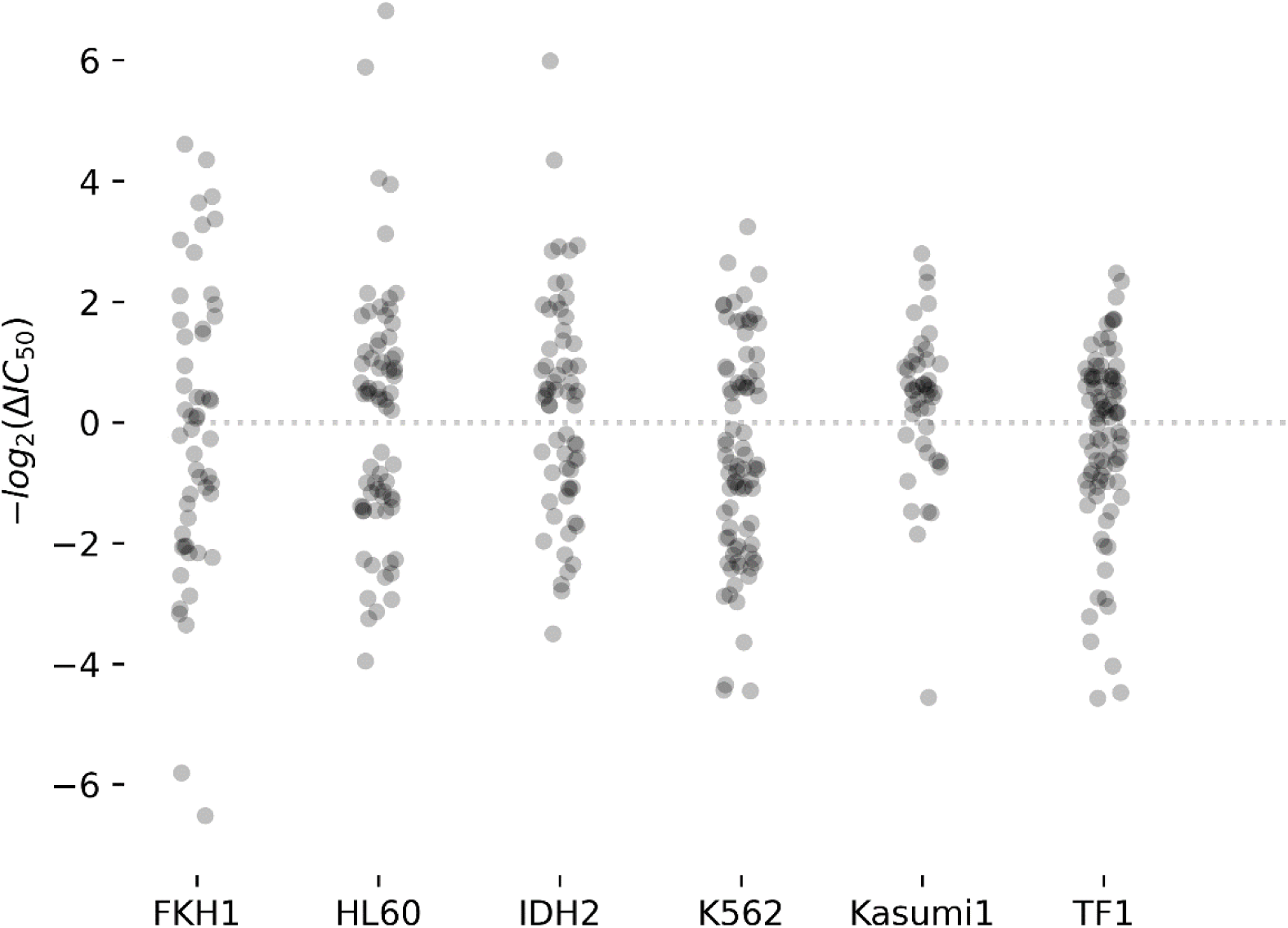
Distribution of synergy scores across all cell lines. Each dot in the scatter plot represents a single drug-drug interaction.

**Supplementary Figure 8.**
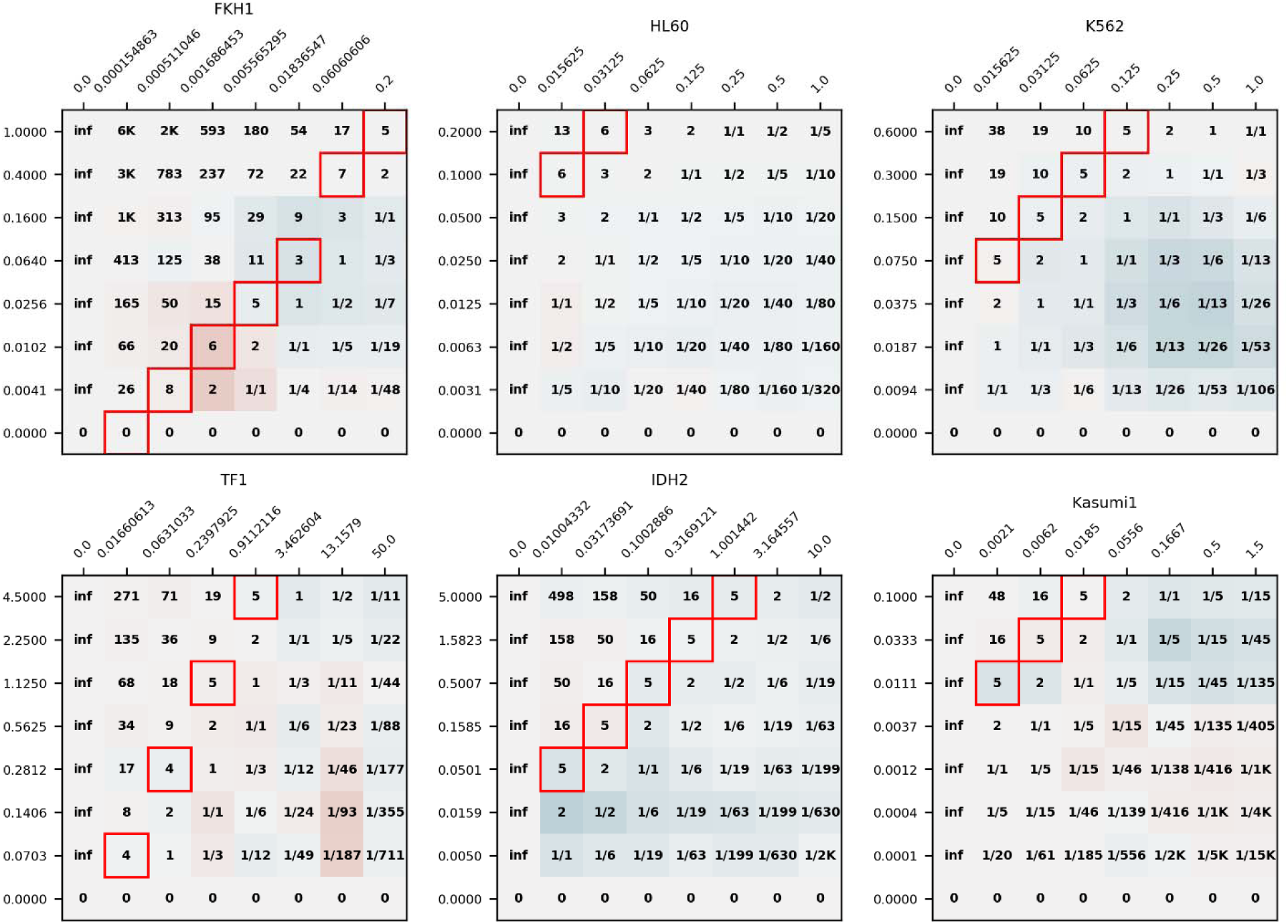
Synergy scores at each concentration level for AraC-DAU. X-axis shows drug concentrations in micromolar for DAU, and y-axis for the AraC. Numbers at the center of each square represents the AraC-to-DAU ratio. Colors indicate the evaluated synergy score against the Bliss independence model. Clinically used typical 5:1 ratio is emphasized with red frames.

**Supplementary Table 1A.** Estimated IC50 values for single drug response curves. Two parameter sigmoid model fits were used (see Methods). Estimation is not available where either a sigmoid model fit was not successful or sufficient killing was not achieved.

|  | FKH1 | HL60 | IDH2 | K562 | Kasumi1 | TF1 |
| --- | --- | --- | --- | --- | --- | --- |
| **2CdA** |  | 0.0225 | 0.1353 | 2.1789 |  | 5.3622 |
| **6MP** |  |  |  |  |  | 3.7643 |
| **6TG** | 11.2732 | 5.6226 | 5.9714 | 1.3195 |  | 2.1266 |
| **ABT199** | 0.0486 | 0.1787 | 10.5327 | 12.0928 | 21.5148 | 13.4339 |
| **AG221** | 62.0053 | 19.9061 | 29.5641 | 41.0308 | 31.5041 | 30.2875 |
| **ATRA** |  | 0.0214 |  | 43.4813 | 205.0733 |  |
| **AZC** | 22.8865 | 3.5871 | 18.4026 | 6.5563 | 6.2519 | 23.9324 |
| **AraC** | 0.0064 | 0.0962 | 0.8647 | 1.0247 | 0.2177 | 2.0250 |
| **DAC** | 18.0545 | 1.9614 | 10.1176 | 6.8441 | 6.3563 | 30.3312 |
| **DAU** | 0.0195 | 0.0057 | 0.2544 | 1.4303 | 0.0241 | 0.3235 |
| **FLD** | 0.2719 | 2.6875 | 11.1980 | 1275.9421 | 10.2013 | 0.0496 |
| **IDA** | 0.0224 | 0.0022 | 0.0871 | 0.0104 | 0.0034 | 0.0399 |
| **MTX** |  |  |  | 0.0101 | 0.0497 | 0.0113 |
| **Mito** | 0.0058 | 0.0030 | 0.1456 | 0.0691 | 0.0152 | 0.1775 |
| **VP16** | 1.0632 | 0.3104 | 10.3568 | 2.5769 | 0.5884 | 9.1563 |

**Supplementary Table 1B.** Estimated b_shape_ coefficients for single drug response curves. Two parameter sigmoid model fits were used (see Methods). Estimation is not available where either a sigmoid model fit was not successful or sufficient killing was not achieved.

|  | **FKH1** | **HL60** | **IDH2** | **K562** | **Kasumi1** | **TF1** |
| --- | --- | --- | --- | --- | --- | --- |
| **2CdA** |  | -3.449 | -0.684 | -0.416 |  | -1.122 |
| **6MP** |  |  |  |  |  | -0.493 |
| **6TG** | -1.186 | -2.738 | -0.816 | -0.704 |  | -0.643 |
| **ABT199** | -0.769 | -0.429 | -3.984 | -1.712 | -1.422 | -2.868 |
| **AG221** | -1.242 | -2.227 | -0.632 | -1.848 | -2.179 | -0.901 |
| **ATRA** |  | -0.141 |  | -1.027 | -0.474 |  |
| **AZC** | -1.088 | -2.029 | -1.699 | -0.397 | -0.645 | -0.899 |
| **AraC** | -0.844 | -0.922 | -0.313 | -0.531 | -0.527 | -0.716 |
| **DAC** | -0.434 | -0.386 | -0.543 | -0.211 | -0.340 | -1.169 |
| **DAU** | -1.009 | -1.634 | -1.203 | -0.688 | -0.677 | -0.982 |
| **FLD** | -5.479 | -2.867 | -1.029 | -0.838 | -3.271 | -1.952 |
| **IDA** | -0.618 | -2.074 | -1.215 | -0.622 | -1.350 | -0.899 |
| **MTX** |  |  |  | -5.495 | -0.299 | -2.786 |
| **Mito** | -1.014 | -1.142 | -0.493 | -0.631 | -1.118 | -0.517 |
| **VP16** | -0.751 | -1.195 | -0.507 | -0.614 | -0.913 | -0.542 |

## Notes

### Competing Interest Statement

The authors have declared no competing interest.

### Summary of Updates

Title revised;Author info updated;Various typing errors fixed;

