## Supplementary Table 4 for "Synergistic and antagonistic drug interactions are prevalent but not conserved across acute myeloid leukemia cell lines"

| Drug name | Drug abbr. | Mechanism of action | Company-Part# | MW (gr/mole) | Plasma min conc. (uM) | Plasma max. conc. (uM) | Reference |
| --- | --- | --- | --- | --- | --- | --- | --- |
| Cytarabine (cytosine arabinoside) | ARA-C | Cytosine analog | Sigma Aldrich C1768 | 243.22 | 82.23 | 242.578 | [1] |
| Azacitidine | AZC | Cytidine analog | Adooq A10105 | 244.2 | 3.071 | 11.261 | [2] |
| Decitabine | DAC | Deoxy form of cytidine analog | Sigma Millipore A3656 | 228.2 | 0.044 | 0.337 | [3] |
| Cladribine | 2CDA | Adenosine analog | Selleck Chem HY-13599 | 285.69 | 10 | 268 | [4] |
| Fludarabine | FLU | 2-Fluoro Ara-A | Selleck Chem S1491 | 285.23 | 0.697 | 3.085 | [5] |
| 6-thioguanine | 6TG | Guanine analog | Sigma Aldrich A4882 | 167.2 | 0.03 | 5 | [6] |
| 6-mercaptopurine | 6MP | Hypoxanthine /guanine analog | Adooq A15898 | 170.19 | 0.3 | 1.8 | [7] |
| Methotrexate | MTX | Folic acid analog | Sigma Aldrich A6770 | 454.44 | 0.1 | 10 | [8] |
| Etoposide | VP16 | Topoisomerase ii inhibitor | Selleck Chem S1225 | 588.56 | 88.011 | 197.941 | [9] |
| Mitoxantrone | MITO | Topoisomerase ii inhibitor | MedChemExpress HY-13599 | 444.48 | 0.058 | 1.0237 | [10] |
| Daunorubicin | DAU | Topoisomerase ii inhibitor | MedChemExpress HY-13062A | 527.52 | 0.01 | 0.032 | [11] |
| idarubicin | IDA | Topoisomerase ii inhibitor | MedChemExpress HY-17381 | 533.95 | 0.019 | 0.129 | [12] |
| all-trans-retinoic acid | ATRA | Differentiation agent | Sigma Millipore R2625 | 300.4 | 119 | 171 | [13] |
| Enasidenib | AG221 | IDH2 R140 inhibitor | MedChem Express  S1225 | 473.38 | 0.858 | 4.289 | [14] |
| Venetoclax | ABT199 | BCL2 inhibitor | MedChem Express S1225 | 868.44 | 0.035 | 4.605 | [15] |

[1] https://doi.org/10.1007/BF02228887

[2] https://www.accessdata.fda.gov/drugsatfda_docs/label/2007/050794s005lbl.pdf

[3] https://doi.org/10.1634/theoncologist.2012-0465

[4] https://doi.org/10.2165/00003088-200342050-00002

[5] https://doi.org/10.1007/BF00256699

[6] https://doi.org/10.1016/B0-12-227555-1/00194-5

[7] https://doi.org/10.1016/B0-12-227555-1/00194-5

[8] https://doi.org/10.22541/au.164866504.46536405/v1

[9] https://doi.org/10.1186/s40780-016-0052-9

[10] https://www.accessdata.fda.gov/drugsatfda_docs/label/2009/019297s030s031lbl.pdf

[11] https://doi.org/10.1016/0304-3835(80)90016-6

[12] https://doi.org/10.1007/BF00686301

[13] https://doi.org/10.1200/JCO.1995.13.5.1238

[14] https://doi.org/10.1182/blood.V126.23.2509.2509

[15] https://doi.org/10.3390/molecules27051607
