## Supplementary Table 5 for "Synergistic and antagonistic drug interactions are prevalent but not conserved across acute myeloid leukemia cell lines"

| **Cell line** | **Origin** | **Company** | **Features** | **Genetics** |
| --- | --- | --- | --- | --- |
| Kasumi-1 (Asou et al., 1991) | 7 yo, M, Japanese  Primary culture from 2nd relapsed BM after BMT | ATCC CRL-2724 | First AML cell line with t(8;21) [a] | t(8,21) |
| HL60 (Collins et al.,1977) | 36 yo F, Caucasian  Primary culture was obtained by leukopheresis prior to treatment. | ATCC CCL-240 | First described as FAB M3 but re-evaluated as FAB M2 [d] erythroleukemia presented like soft tissue myeloid tumors [b] | c-myc  t(5;17) and der(16)t(5;16) |
| FKH1 (Hamaguchi et al., 1998) | 61 yo M, Japanese  peripheral blood was obtained after CML to AML transformation. | DSMZ  ACC 614 | AML M4 was transformed from refractory  Ph(-) CML with trilineage myelodysplasia | t(6;9)(p23;q34)  leading to  DEK-NUP214  (DEK-CAN) fusion gene |
| TF1 (T. Kitamura, et al.,1987) | 35 yo M, Japanese  Cells were obtained from BMA sample of severe pancytopenic patient. | ATCC  CRL-2003 | Erythroleukemia |  |
| IDH2 mutated TF1 | 35 yo M, Japanese | ATCC  CRL-2003IG | Erythroleukemia | (AML) IDH2R140Q mutant isogenic line derived from the parental TF-1 cell line by using CRISPR/Cas9 technology followed by single cell cloning |
| K562 (Lozzio and Lozzio, 1975) | 53 yo F  Cells were obtained from pleural effusion of R/R CML | ATCC  CCL-243 | Erythroleukemia | t(15;17)(q21;q24) Ph1:deI(22)(q12) NOTCH1  E2A GATA2  GATA1 |
